# Molecular and clinicopathological characterization of a prognostic immune gene signature associated with MGMT methylation in glioblastoma

**DOI:** 10.1101/2020.07.16.206318

**Authors:** Liang Zhao, Jiayue Zhang, Shurui Xuan, Zhiyuan Liu, Yu Wang, Peng Zhao

**Author notes:** **Correspondence to:** Peng Zhao, Department of Neurosurgery, The First Affiliated Hospital of Nanjing Medical University, Nanjing, Jiangsu, 210000, China. These authors contributed equally to this work (Liang Zhao, Jiayue Zhang).

## Abstract

**Background:** O6-methylguanine-DNA methyltransferase (MGMT) methylation status affects tumor chemo-resistance and the prognosis of glioblastoma (GBM) patients. We aimed to investigate the role of MGMT methylation in the regulation of GBM immunophenotype and discover an effective biomarker to improve prognosis prediction of GBM patients.

**Methods:** A total of 769 GBM patients with clinical information from five independent cohorts were enrolled in the present study. Samples from the Cancer Genome Atlas (TCGA) dataset were used as the training set, whereas transcriptome data from the Chinese Glioma Genome Atlas (CGGA) RNA-seq, CGGA microarray, GSE16011, and the Repository for Molecular Brain Neoplasia (REMBRANDT) cohort were used for validation. A series of bioinformatics approaches were carried out to construct a prognostic signature based on immune-related genes, which were tightly related with the MGMT methylation status. The influence of the signature on immunosuppression and remodeling of the tumor microenvironment were comprehensively investigated. Then, the utility of this immune gene signature was analyzed by the development and evaluation of a nomogram.

**Results:** We found that MGMT unmethylation was closely associated with immune-related biological processes in GBM. Sixty-five immune genes were more highly expressed in the MGMT unmethylated than the MGMT methylated group. An immune gene-based risk model was further established to divide patients into high and low-risk groups, and the prognostic value of this signature was validated in several GBM cohorts. Functional analyses manifested a universal up-regulation of immune-related pathways in the high-risk group as compared to the low-risk group. Furthermore, the risk score was highly correlated to the immune cell infiltration, immunosuppression, inflammatory activities, as well as the expression levels of immune checkpoints. Finally, a nomogram was developed for clinical application.

**Conclusions:** MGMT methylation is strongly related to the immune responses in GBM. The immune gene-based signature we identified may have potential implications in predicting the prognosis of GBM patients and mechanisms underlying the role of MGMT methylation.

## Background

Glioma is a type of central nervous system (CNS) neoplasms and accounts for the majority of intracranial malignant tumors in adults. Glioblastoma (GBM), which is defined as the World Health Organization (WHO) grade IV glioma, shows a highly aggressive, heterogeneous, and lethal phenotype. Currently, the etiology of GBM is still largely unknown. Although surgical resection combined with chemotherapy or radiotherapy has been widely used as a routine clinical treatment, the prognosis of patients has even not improved significantly, with a median overall survival fewer than 15 months (1). Due to the persistent proliferation of tumor cells and penetration into the surrounding tissues, total tumor resection is seldom possible (2). Alkylating agents, such as temozolomide(TMZ)andprocarbazine, are appliedas first-line chemotherapeutic drugs (3). However, many GBM patients do not respond to alkylating agents or develop chemo-resistance after a period of chemotherapy (4). Therefore, there is an urgent need to explore the underlying mechanisms of gliomagenesis and develop reliable therapeutic approaches.

The intracellular DNA repair enzyme O^6^-methylguanine-DNA methyltransferase (MGMT) catalyzes the DNA repair process by removing the alkylation of the O^6^ position of guanine (5). By generating methylguanine adducts, DNA-alkylating drugs can trigger base mismatching during DNA replication, affecting the cell cycle and promoting the death of tumor cells (6) (7) (8). As a most striking modification form, methylation of the MGMT promoter can induce a decrease of MGMT expression and further improve the curative effect of chemotherapy with alkylating agents (1). In the case of GBM, clinical trials revealed that patients with MGMT methylation benefit more than those with unmethylated tumors when received combined radiochemotherapy (9). Furthermore, patients without MGMT methylation have a shorter overall survival time (10). So far, these underlying molecular mechanisms have not been exhaustively described, especially about tumor immune microenvironment.

Recently, breakthroughs in tumor immunotherapy have revolutionized the treatment of several cancer types (11). The specific intratumoral immune microenvironment can facilitate immune escape, drive tumorigenesis, and promote the malignant progression of the lesion (12) (13). Immune checkpoints play an intermediary role in the tumor-immune system dynamics, and inhibitory immune checkpoints have been studied intensively as therapeutic targets for multiple malignant tumors. In non-small cell lung cancer (NSCLC) and melanoma, the application of immunosuppressive agents has produced remarkable antitumor effects (14) (15). The discovery of lymphatic system vessels in the CNS offered new hope for developing novel immunotherapeutic methods for GBM (16). The first large phase III clinical trial of nivolumab combined with ipilimumab (PD-1 inhibitors) in recurrent GBM (NCT02017717) was launched in 2014. A recent clinical trial indicated that the neoadjuvant *PD-1* blockade could upregulate the amount of T cells, promote the expression of interferon-γ-related genes, and further offer a promising survival benefit in recurrent GBM (17). The previous study has demonstrated that isocitrate dehydrogenase (IDH) mutation status was closely associated with immune response in GBM, and a prognostic model based on IDH mutation was further developed to predict the clinical outcomes of patients (18). However, the role of MGMT methylation status in regulating tumor immunity in GBM remains to be discussed, and an MGMT methylation-related biomarker should be exploited.

Here, we utilized data from a large number of GBM samples from multiple publicly available datasets to comprehensively investigate the potential relationship between MGMT status and the immunological tumor microenvironment. We hypothesized that the shorter survival of patients with MGMT-unmethylated GBM is partly due to the pro-tumor immune response. More importantly, we developed a robust prognostic model based on MGMT methylation status to predict the clinical outcomes of GBM patients.

## Methods

### GBM patient datasets

We retrospectively collected a large-scale profile composed of five independent datasets involving 769 GBM patients (Table S1). The RNA sequencing (RNA-seq) gene expression data for 165 GBM samples were obtained from The Cancer Genome Atlas (TCGA) database. The microarray and RNA-seq transcriptome data for 112 and 113 GBM patients were downloaded from the Chinese Glioma Genome Atlas (CGGA) database (www.cgga.org.cn). The microarray gene expression profile matrix files of the GSE16011 project (including 159 GBM samples) and the Repository for Molecular Brain Neoplasia (REMBRANDT) database (including 220 GBM samples) were downloaded from the Gene Expression Omnibus (GEO) database (www.ncbi.nlm.nih.gov/geo/). The corresponding profiles comprising detailed clinicopathological characteristics were also generated from each data source. The TCGA RNA-seq dataset was used as the training set and the CGGA RNA-seq dataset as the validation set. Furthermore, the CGGA microarray profile, as well as the GSE16011 and REMBRANDT datasets, were used to test the prognostic value of the risk model developed in the TCGA cohort.

### Immune-related genes

The 1039 known immunologically relevant genes were downloaded from the ImmPort database (www.immport.org), and 1100 immune-related genes were extracted from the “IMMUNE_RESPONSE” gene set (Molecular Signatures Database V7.0, http://software.broadinstitute.org/gsea/msigdb/index.jsp). Finally, after combining these two independent gene sets, 1763 immune-related genes were selected for the next analyses (Table S2).

### Identification of prognostic MGMT methylation related immune genes

137 GBM patients with data on gene expression and MGMT methylation status was selected in the TCGA cohort, and the differentially expressed genes (DEGs) between MGMT unmethylated and methylated samples were identified using the “DESeq2” R package (19). Before performing the differential gene expression analysis, low-abundance genes with raw counts less than 10 in more than 75% of samples were removed (20). Then, the differentially expressed immune genes were filtered using the criterion of log2 fold change > 1 and FDR-adjusted P < 0.05. Similarly, DEGs (log2 fold change > 0.8 and adjusted P < 0.05) between different MGMT methylation status in the CGGA RNA-seq cohort were identified using the “limma” R package (21). The prognostic value of the identified genes in the TCGA cohort was further investigated using a multivariate Cox regression model integrating prognostic clinicopathological information, including age, Karnofsky performance score (KPS), IDH mutation, and MGMT methylation status. Then, genes with significant prognostic value (P < 0.05) were screened for further analyses.

### Construction of a risk model based on immune-related genes

The least absolute shrinkage and selection operator (LASSO) regression model with 10-fold cross-validation was adopted to select the most useful prognostic factors for GBM patients. The optimal lambda value was estimated based on the minimum criteria. Then, the risk score for each sample was calculated using the expression levels of genes and corresponding regression coefficients obtained from the multivariable Cox regression analysis. The formula of the risk score model was as follows:

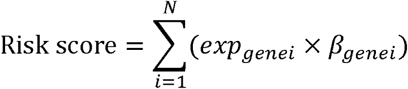

where N is the number of genes in the signature, β_genei_ is the regression coefficient, and exp_genei_ represents the expression value of a specific gene in a single sample. The cutoff for classifying GBM patients into high and low-risk groups was determined using X-title 3.6.1 software (22). Kaplan-Meier survival analysis was used to evaluate the predictive value of the risk model, and time-dependent receiver operating characteristic (ROC) curves were drawn to assess the efficiency of the risk model in predicting the clinical outcomes at different times. This procedure was completed using the “timeROC” R package.

### Functional enrichment analyses

Metascape, an easy-to-use web portal that provides a comprehensive analysis for the functional annotation of lists of genes (23), was used to perform Gene Ontology (GO) term enrichment and the Kyoto Encyclopedia of Genes and Genomes (KEGG) pathway analysis for genes of interest. Gene set enrichment analysis (GSEA) analysis was conducted using the R package “fgsea” (24). GSEA gene set (C5 collection) was downloaded from the Molecular Signatures Database (v7.0). To obtain robust results, we performed gene-set permutations for 10000 times during GSEA. Enriched terms with FDR-adjusted P < 0.05 were deemed as statistically significant.

### Quantification of immune-cell infiltration by single-sample gene set enrichment analysis (ssGSEA)

The estimation of overall immune cell type fractions in GBM samples was performed using the ssGSEA algorithm in the “GSVA” R package (25), which uses 24 different types of immune cell marker genes derived from previous research to analyze each GBM sample (26). Markers associated with cells of the innate immune system, including natural killer (NK) cells, NK CD56^dim^ cells, NK CD56^bright^ cells, dendritic cells (DCs), immature DCs (iDCs), activated DCs (aDCs), neutrophils, mast cells, eosinophils, and macrophages, as well as those associated with cells of the adaptive immune system, including B, T central memory (Tcm), CD8^+^ T, T effector memory (Tem), T follicular helper (Tfh), Tγδ, Th1, Th2, Th17, and Treg cells, were included in the gene list. A numeric matrix consists of the enrichment score of each immune cell type in a single sample was obtained via ssGSEA.

### Prediction of clinical responses to immune checkpoint blockade

We used the computational method of Tumor Immune Dysfunction and Exclusion (TIDE) (27) to predict the likelihood of response to immune checkpoint blockade therapy for GBM patients. Due to a lack of sufficient data from normal brain tissues in the TCGA dataset, the TCGA TARGET GTEx dataset from UCSC Toil RNA-seq Recompute project (28), which includes a conjoint analysis of three independent datasets in order to make the gene expression of samples from different datasets or batches comparable, was adopted for subsequent studies. The gene expression values of GBM patients were normalized towards the corresponding average values of normal brain tissues (cortex, frontal cortex (Ba9), and anterior cingulate cortex (Ba24)) from the GTEx project before subjecting the data to the TIDE algorithm. Next, the subclass mapping method (SubMap) (29) was used to compare the gene expression matrices of different risk groups with the expression profile of melanoma patients who received immunotherapy targeting *CTLA-4* and *PD-1 (30)*. This step was performed using the default parameters in the SubMap module on the GenePattern website (http://genepattern.broadinstitute.org/).

### Construction and validation of a nomogram

Multivariable Cox proportional hazards regression analysis was applied with the following clinical-relevant covariates: gender, age, MGMT methylation status, Karnofsky Performance Status (KPS) score, IDH mutation status, and risk score. A combined nomogram was generated as a quantitative tool for predicting the likelihood to die of each patient using the “regplot” R package. The concordance index (C-index) was calculated to assess the consistency between model prediction and actual clinical outcomes of patients. The calibration plot was generated to evaluate the accuracy of the prediction for 1-, 2-, and 3-year overall survival using this nomogram by the “rms” R package.

### Statistical analysis

The LASSO Cox regression analysis was carried out using the “glmnet” R package. Restricted mean survival (RMS) represents the loss in average life expectancy for patients, and the RMS, as well as time ratio for the risk model in each dataset, were calculated using the “survRM” R package. Tumor purity, which represents the heterogeneity of each tumor sample, was estimated by the “ESTIMATE” R package (31). Univariate and multivariate Cox regression analyses were used to identify the risk-score-based signature as an independent prognostic factor for GBM patients. R software (version 3.6.1, www.r-project.org) was used for all statistical analyses. Fisher’s exact test was used to compare the differences of clinicopathologic features as well as immunotherapy response between high- and low-risk groups, and was implemented using the “fisher.test” function in R software. Mann-Whitney U test was used to assess the statistical significance of differences between the means of continuous data. All hypothetical tests were two-sided, and P values < 0.05 were considered to indicate statistical significance.

## Results

### Association between MGMT methylation and the immunological phenotype of GBM

There is a large body of work implicating that MGMT methylation can affect the prognosis as well as the effectiveness of chemotherapy in GBM. However, the influence of MGMT promoter status on the immunological phenotype of GBM had, to our best knowledge, not been investigated prior to this study. Here, we utilized the RNA-seq gene expression profiles and clinical information of GBM patients from both the TCGA and CGGA RNA-seq databases to explore the molecular differences in underlying immunological mechanisms according to the MGMT methylation status. Patients were divided into MGMT^U^ (MGMT-unmethylated) and MGMT^M^ (MGMT methylated) groups in these two cohorts, respectively. GSEA analysis showed that immune-related biological processes, such as humoral immune response (normalized enrichment score (NES) = 1.95, FDR < 0.001), T cell activation involved in immune response (NES = 1.75, FDR < 0.001), adaptive immune response (NES = 1.72, FDR < 0.001), cytokine production involved in immune response (NES = 1.65, FDR < 0.001), and regulation of innate immune response (NES = 1.30, FDR < 0.01), were significantly enriched in the MGMT^U^ group in the TCGA cohort (Figure 1A). Notably, all immune-related GO terms with NES > 1 were only enriched in the MGMT^U^ group in the TCGA cohort (Table S3). In another independent GBM dataset (CGGA RNA-seq cohort), immune-related biological processes were also highly enriched in the MGMT^U^ samples (Figure 1B, Table S3). Top enriched gene terms were all implicated in regulating immune activities in these two GBM datasets. GO and KEGG analyses showed that MGMT-unmethylation-specific upregulation of genes associated with the immune-related biological processes and signaling pathways (Figure 1C, 1D). Genes with roles in the activation and migration of different immune cells, cytokine and chemokine signaling axes, and IL-17 pathway were characterized by enrichment of genes associated with immunological functions. These findings suggested that the MGMT promoter methylation status may affect the prognosis of GBM patients by mediating the immune response in the malignant tumor environment.

**Figure 1.**
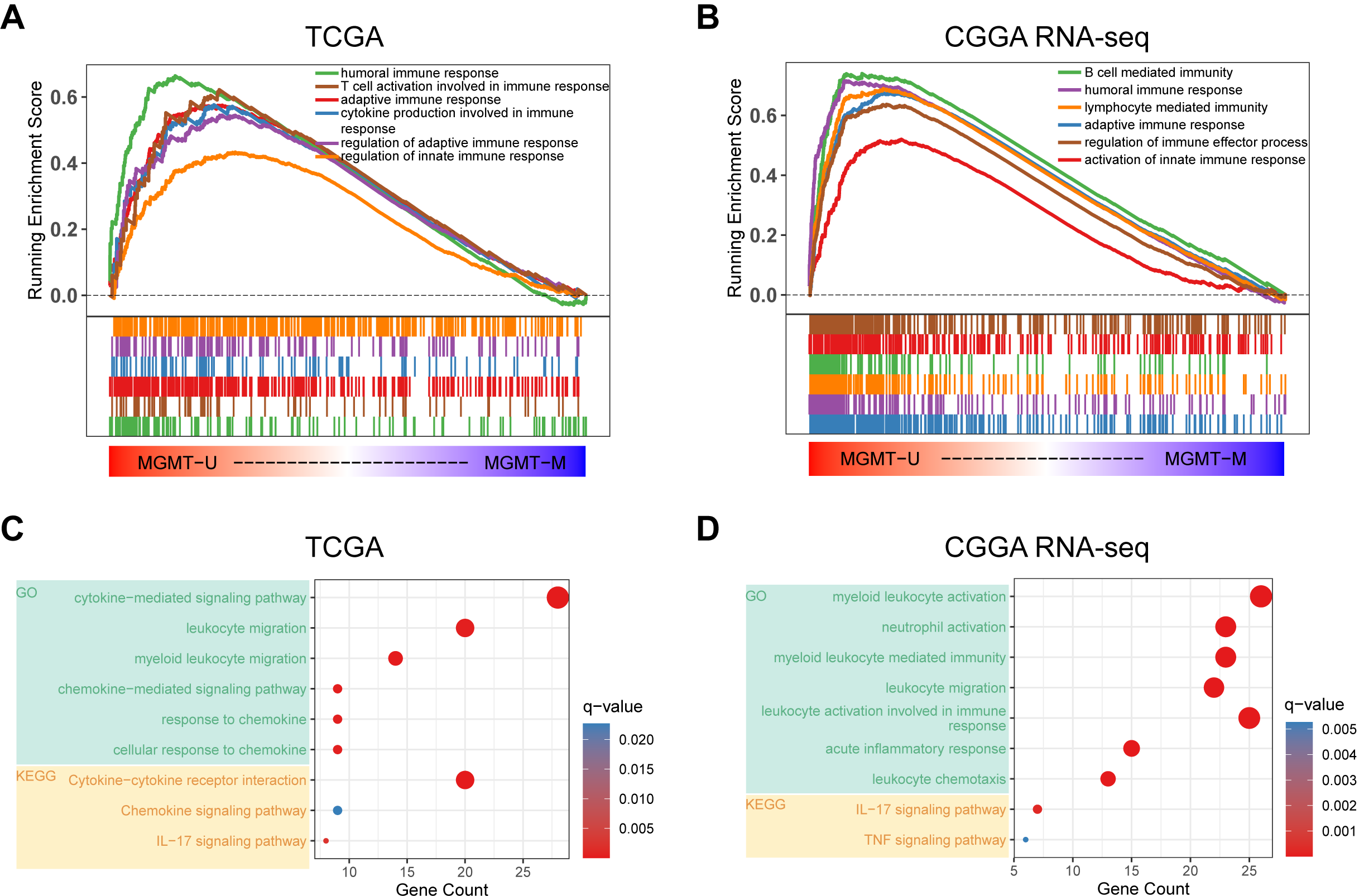
Functional annotation of MGMT methylation status in the TCGA and CGGA RNA-seq datasets. Gene Set Enrichment Analyses showing genes upregulated in MGMT unmethylated samples evaluated in the gene sets representative for immune-related biological processes and pathways in both TCGA (**A**) and CGGA RNA-seq (**B**) GBM cohorts. Normalized enrichment score (NES) and FDR were generated for each term. GO and KEGG analyses showing the enriched biological processes and signaling pathways in MGMT unmethylated samples of the TCGA (**C**) and CGGA RNA-seq (**D**) datasets.

### Construction of an immune risk model based on MGMT methylation status

To further investigate the potential immune pathways and key immune-related genes driving this phenomenon, differentially expressed genes between the MGMT^U^ and MGMT^M^ groups in the TCGA database were identified (Figure 2A, Table S4). Sixty-five immune-related genes were found to be more highly expressed in the MGMT^U^ than the MGMT^M^ group (log2 fold change > 1, FDR-adjusted P < 0.05), such as CXC motif chemokine ligand (CXCL1, 2, 6, 12, 13), immunoglobulin kappa variable family (IGKV1-27, IGKV2-28, IGKV3-11, IGKV3-15), and immunoglobulin heavy constant gamma cluster (IGHG1, 2, 3).

**Figure 2.**
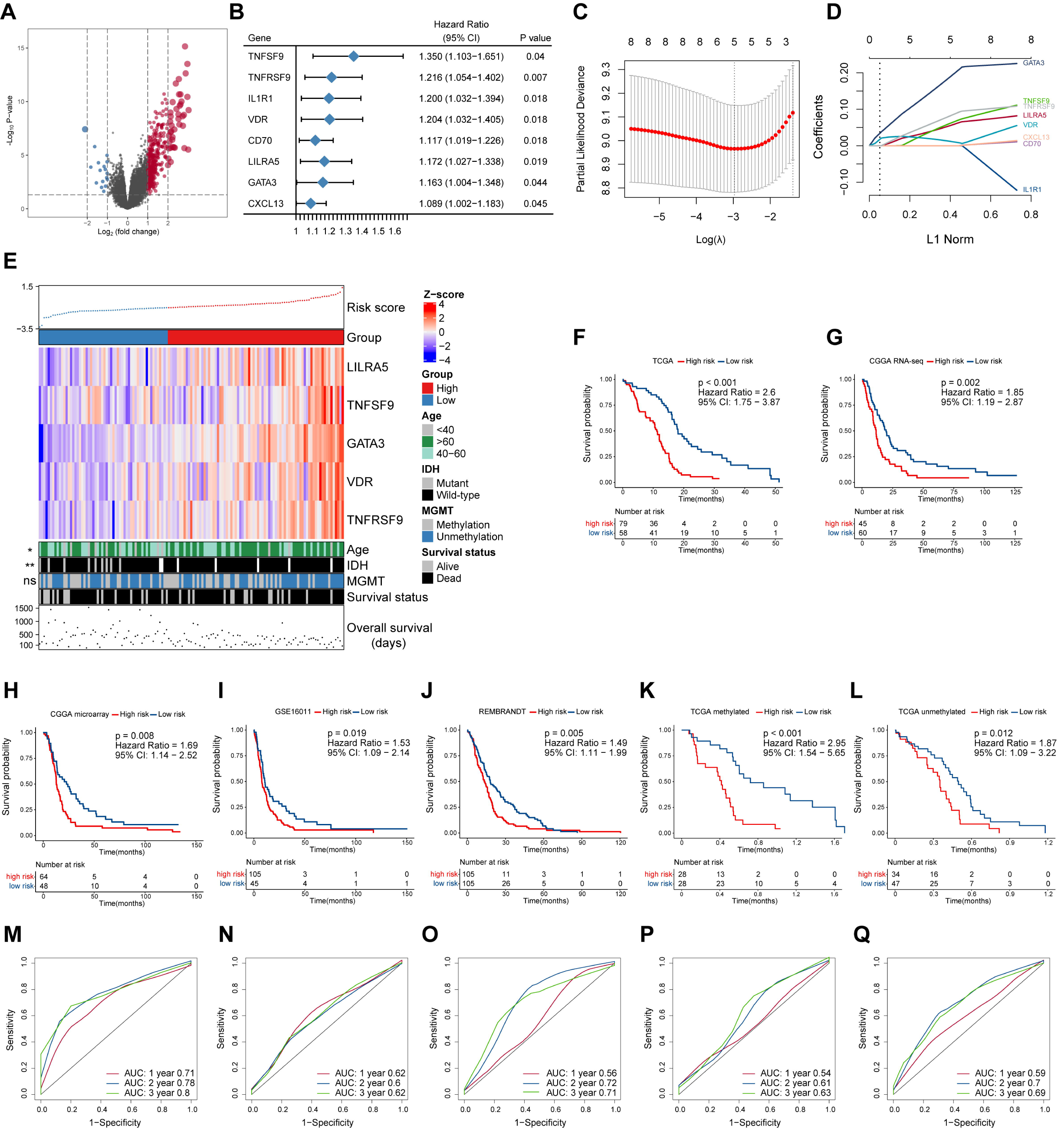
Construction and prognostic analysis of the immune gene-based signature. **A**. Volcano plot of fold changes of genes in groups with different MGMT methylation status in the TCGA dataset. Overexpressed genes in MGMT unmethylated group were plotted as red dots while blue dots represent downregulated genes. The x and y-axis represent fold-change and the P-values estimated by DESeq2. **B**. Results of multivariate Cox regression analysis using differentially expressed immune genes in different MGMT methylation status. Genes with p-values less than 0.05 were obtained. **C**. The partial likelihood deviance was plotted using vertical lines with the red dot, and the dotted vertical lines represent values based on minimum criteria and 1-SE criteria, respectively. **D**. LASSO coefficient profiles of the candidate prognostic immune genes by 10-fold cross-validation. **E**. The relationship between the expression patterns of five candidate immune genes and the increasing risk score, as well as other clinicopathological features. Kaplan-Meier analysis of overall survival based on high *vs*. low risk in different cohorts (**F-J**). The prognostic value of this model was further validated in subgroups with MGMT methylation (**K**) and unmethylation (**L**) from the TCGA dataset. P values were obtained from the log-rank test. Time-dependent ROC curves analysis of this signature in the TCGA (**M**), CGGA RNA-seq (**N**), CGGA microarray (**O**), GSE16011 (**P**), and REMBRANDT (**Q**) datasets.

The prognostic effect of the differentially expressed immune genes was investigated by applying multivariate Cox regression adjusted for other survival-related clinicopathological covariates. TNF superfamily member 9 (TNFSF9), TNF receptor superfamily member 9 (TNFRSF9), Interleukin 1 receptor type 1 (IL1R1), vitamin D receptor (VDR), CD70, leukocyte immunoglobulin like receptor A5 (LILRA5), GATA binding protein 3 (GATA3), and CXC motif chemokine ligand 13 (CXCL13) were selected as independent factors associated with the overall survival of GBM patients (Figure 2B, Table S5). All these immune genes were regarded as risk factors with a hazard ratio (HR) > 1. Among these candidates, LASSO Cox regression identified five highly relevant immune-related genes: LILRA5, TNFSF9, GATA3, VDR, and TNFRSF9 (Figure 2C, 2D). The robustness of the result was confirmed using 10-fold cross-validation. Circos plots (32) were generated to delineate the correlations between these genes in both the TCGA and CGGA RNA-seq datasets (Figure S1A, 1B). It is worth noting that the relationships of these genes were quite similar in different datasets indicating that the identified relevant immune genes had a stable positive correlation in GBM.

A schematic view of MGMT methylation related immune gene selection and prognostic gene signature development is delineated in Figure S2. The risk score of each GBM patient was calculated by combining the expression level of genes and corresponding coefficients derived from the multivariate Cox regression analysis. Patients were split into high- and low-risk populations according to the optimal cutoff value estimated using X-title software. The relationship between the expression patterns of these five prognostic immune genes and the increasing risk score in the TCGA cohort were shown (Figure 2E). More older and IDH mutated patients were found in the high-risk group compared with those with low risk score, while no difference in the proportion of MGMT methylation between these groups was found. Kaplan–Meier survival analysis further showed that patients from the high-risk group had shorter overall survival than those with low risk in the TCGA cohort (HR = 2.6, 95% confidence interval (CI): 1.75-3.87, P < 0.001) (Figure 2F). We also applied this computational formula to the CGGA RNA-seq, CGGA microarray, GSE16011, and REMBRANDT datasets to explore the prognostic value of the identified immune-genetic signature. Consistent with the results of the TCGA cohort, high-risk patients had significantly worse outcomes than those with low risk (Figure 2G-2J). Significant RMS time ratios ranging from 1.465 to 1.854 were observed in the five datasets (P < 0.05, Table S6). In the TCGA dataset, we also checked the efficiency of this model in patients with or without MGMT methylation (Figure 2K, 2L). The results showed that low-risk patients had a significant survival advantage over the high-risk group in these two counterparts (P < 0.05). Time-dependent ROC curves were generated to evaluate the efficiency of this risk signature in predicting 1-, 2-, and 3-year survival of GBM patients, and the risk model showed considerable predictive potential in different cohorts (Figure 2M-2Q).

To confirm whether the risk model can be used as an independent prognostic tool for GBM, univariate and multivariate Cox regression analyses were applied to the TCGA dataset. The risk score signature was adjusted using prognostic information of clinical factors that had been deemed statistically significant in univariate analysis (P < 0.05). The HRs for the signature in the univariate and multivariate analyses were 3.107 (P < 0.001, 95% CI: 2.005-4.816) and 2.476 (P < 0.001, 95% CI: 1.581-3.878), respectively. The risk model was further validated in the CGGA RNA-seq cohort, with HRs of 1.896 (P = 0.003, 95% CI: 1.246-2.886) and 1.644 (P = 0.03, 95% CI: 1.05-2.573), respectively (Table 1). Thus, the signature based on immune-related genes can be used independently to predict the overall survival of GBM patients.

**Table 1.**
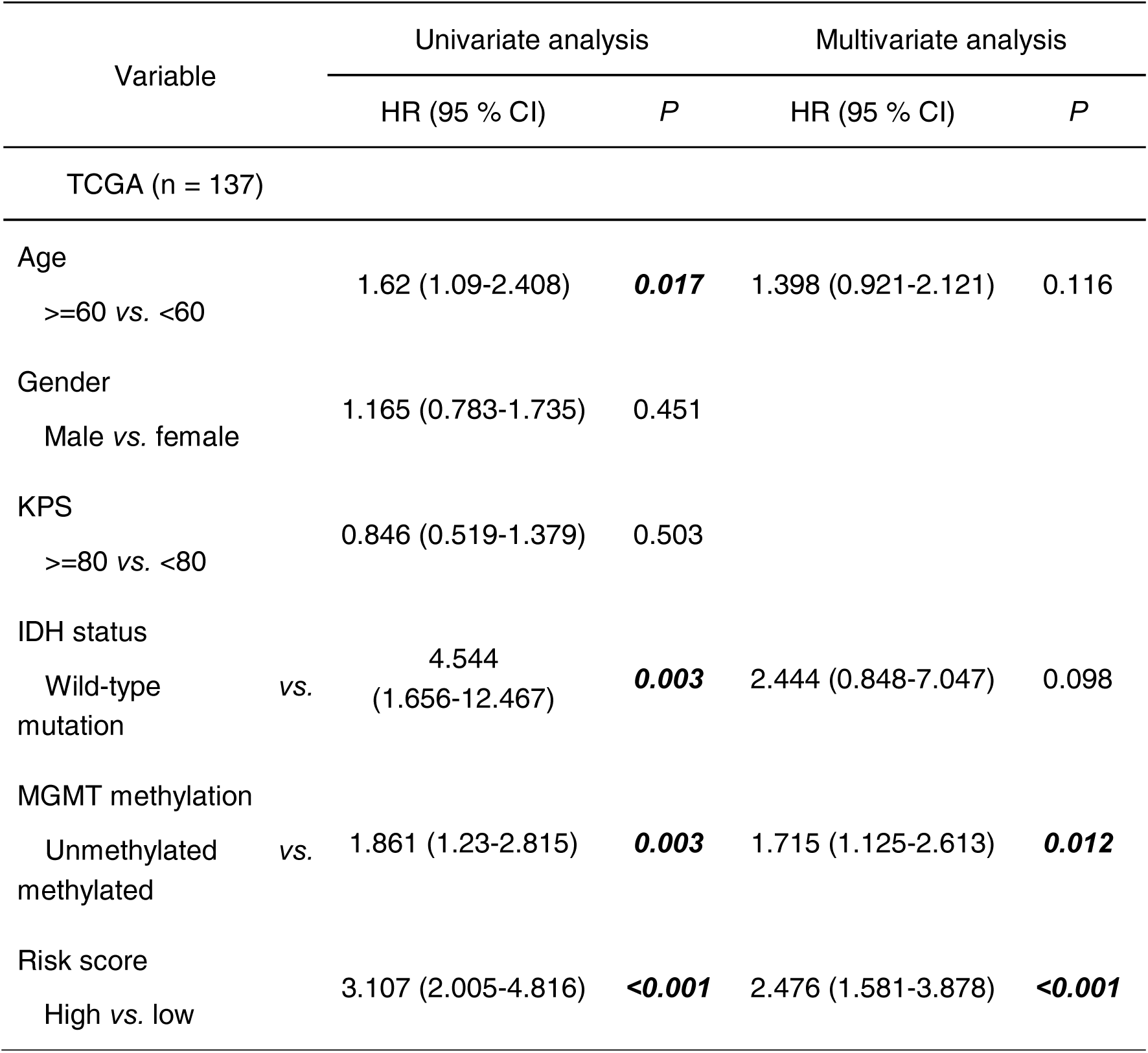
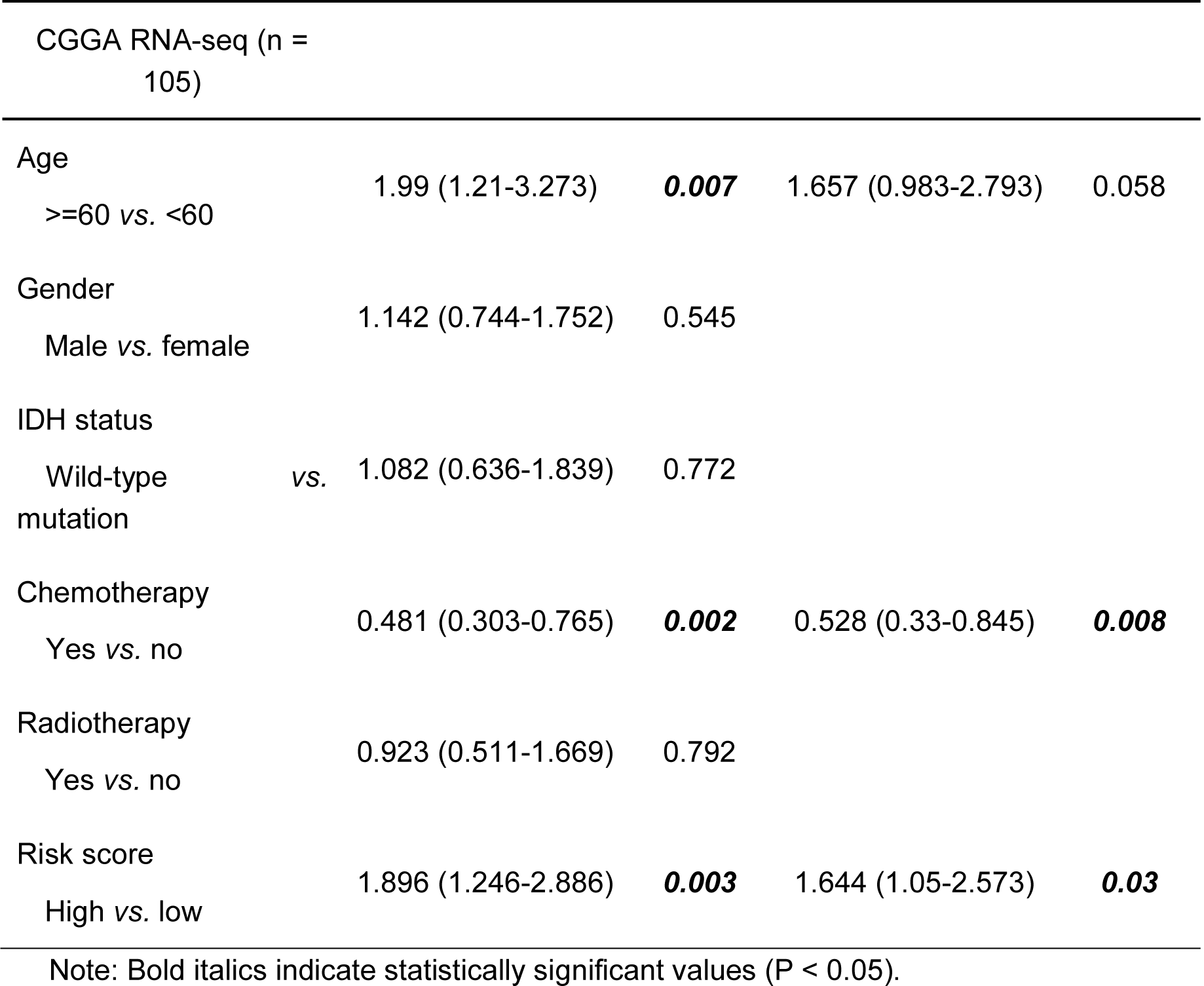
Univariate and multivariate analyses of clinicopathological characteristics and immune gene-based signature with overall survival in GBM cohorts.

| Variable | Univariate analysis |  | Multivariate analysis |  |
| --- | --- | --- | --- | --- |
|  | HR (95 % CI) | *P* | HR (95 % CI) | *P* |
| TCGA (n = 137) |  |  |  |  |
| Age |  |  |  |  |
| >=60 vs. <60 | 1.62 (1.09-2.408) | **0.017** | 1.398 (0.921-2.121) | 0.116 |
| Gender |  |  |  |  |
| Male vs. female | 1.165 (0.783-1.735) | 0.451 |  |  |
| KPS |  |  |  |  |
| >=80 vs. <80 | 0.846 (0.519-1.379) | 0.503 |  |  |
| IDH status |  |  |  |  |
| Wild-type vs. mutation | 4.544<br>(1.656-12.467) | **0.003** | 2.444 (0.848-7.047) | 0.098 |
| MGMT methylation |  |  |  |  |
| Unmethylated vs. methylated | 1.861 (1.23-2.815) | **0.003** | 1.715 (1.125-2.613) | **0.012** |
| Risk score |  |  |  |  |
| High vs. low | 3.107 (2.005-4.816) | **<0.001** | 2.476 (1.581-3.878) | **<0.001** |

| CGGA RNA-seq (n = 105) |  |  |  |  |
| --- | --- | --- | --- | --- |
| Age |  |  |  |  |
| ≥60 vs. <60 | 1.99 (1.21-3.273) | **0.007** | 1.657 (0.983-2.793) | 0.058 |
| Gender |  |  |  |  |
| Male vs. female | 1.142 (0.744-1.752) | 0.545 |  |  |
| IDH status |  |  |  |  |
| Wild-type vs. mutation | 1.082 (0.636-1.839) | 0.772 |  |  |
| Chemotherapy |  |  |  |  |
| Yes vs. no | 0.481 (0.303-0.765) | **0.002** | 0.528 (0.33-0.845) | **0.008** |
| Radiotherapy |  |  |  |  |
| Yes vs. no | 0.923 (0.511-1.669) | 0.792 |  |  |
| Risk score |  |  |  |  |
| High vs. low | 1.896 (1.246-2.886) | **0.003** | 1.644 (1.05-2.573) | **0.03** |
| Note: Bold italics indicate statistically significant values (P < 0.05). |  |  |  |  |

### Functional analysis of the MGMT methylation-based signature

To further identify the potential biological processes and signaling pathways associated with the immune gene signature, GO and KEGG functional analyses were performed, and the results were graphed using Metascape online tool. Firstly, we selected a list of genes that were strongly correlated with the risk score (|Pearson correlation coefficient| (|R|) > 0.4, P < 0.05) in the TCGA and CGGA RNA-seq cohorts (Table S7). Heatmaps for these genes and corresponding detailed clinicopathological features were plotted, as depicted in Figure S3A, 3B. Functional analysis demonstrated that these genes were significantly enriched in immune processes, such as lymphocyte activation, cytokine production, cytokine-mediated signaling pathway, leukocyte migration, activation of the immune response, regulation of the inflammatory response, and interferon-gamma production (P < 0.05) (Figure 3A, 3B). Similarly, immune-related terms such as cytokine-mediated signaling pathway, myeloid leukocyte activation, cytokine production, leukocyte migration, TNF signaling pathway, and NF-kappa B signaling pathway were also enriched in genes derived from the CGGA RNA-seq cohort (P < 0.05) (Figure S4A, 4B).

**Figure 3.**
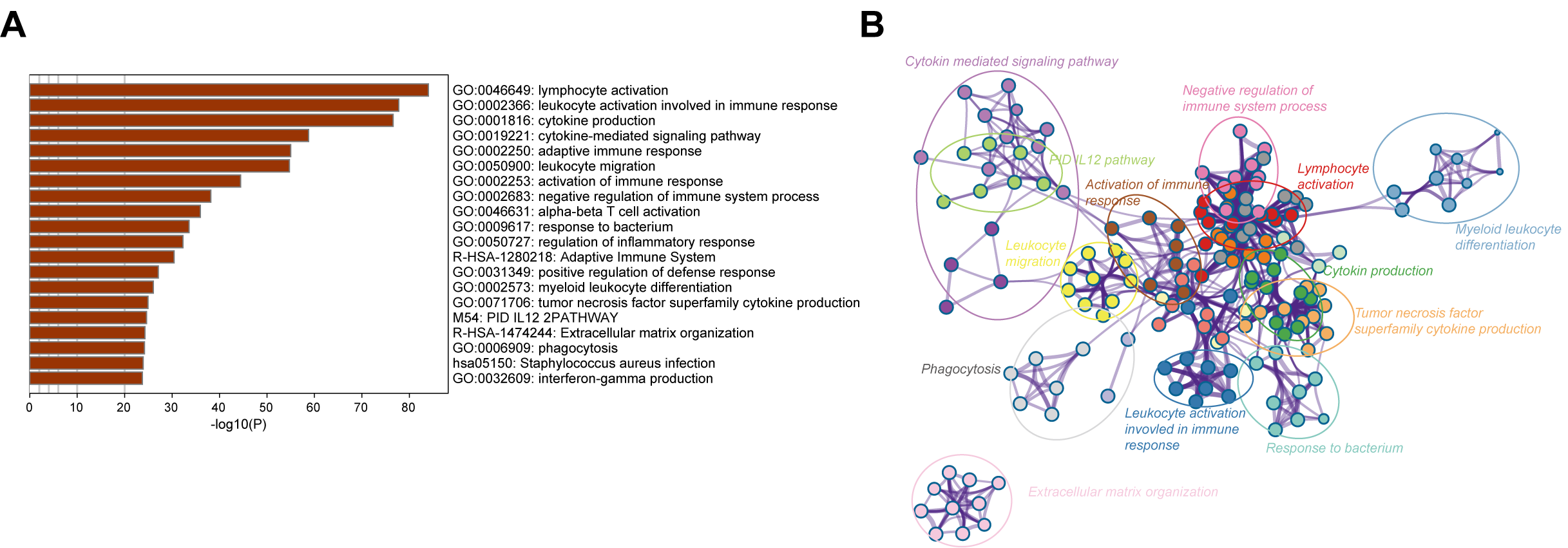
Biological functions of the immune gene signature in GBM in the TCGA dataset. **A**. Bar plot showing the top 20 terms derived from the gene set enrichment of risk score. The x-axis represents statistical significance. **B**. The enrichment network plot visualizing the relationship between a set of representative terms. Each term is assigned with a unique color.

### Association of the risk score with tumor purity and immune cell infiltration in the GBM microenvironment

The intratumoral niche is a complex microenvironment that consists of the tumor and non-tumor cells, such as stromal and immune cells (33). It is now clear that a mass of non-tumor cells plays an important role in the initiation and progression of various human cancers (34) (35) (36). Tumor purity is defined as the proportion of cancer cells in the mixed tumor tissues (37) and has been identified as an effective prognostic index in glioma (2). To clarify the relationship between the immune gene-based risk score and the tumor purity, we used the ESTIMATE algorithm to calculate the purity of GBM samples based on the transcriptomic gene expression data. In both the TCGA and CGGA RNA-seq cohorts, the risk score was negatively correlated with the tumor purity (R = −0.6, P < 0.0001 in TCGA, and R = −0.4, P <0.0001 in CGGA RNA-seq) (Figure 4A, 4B).

**Figure 4.**
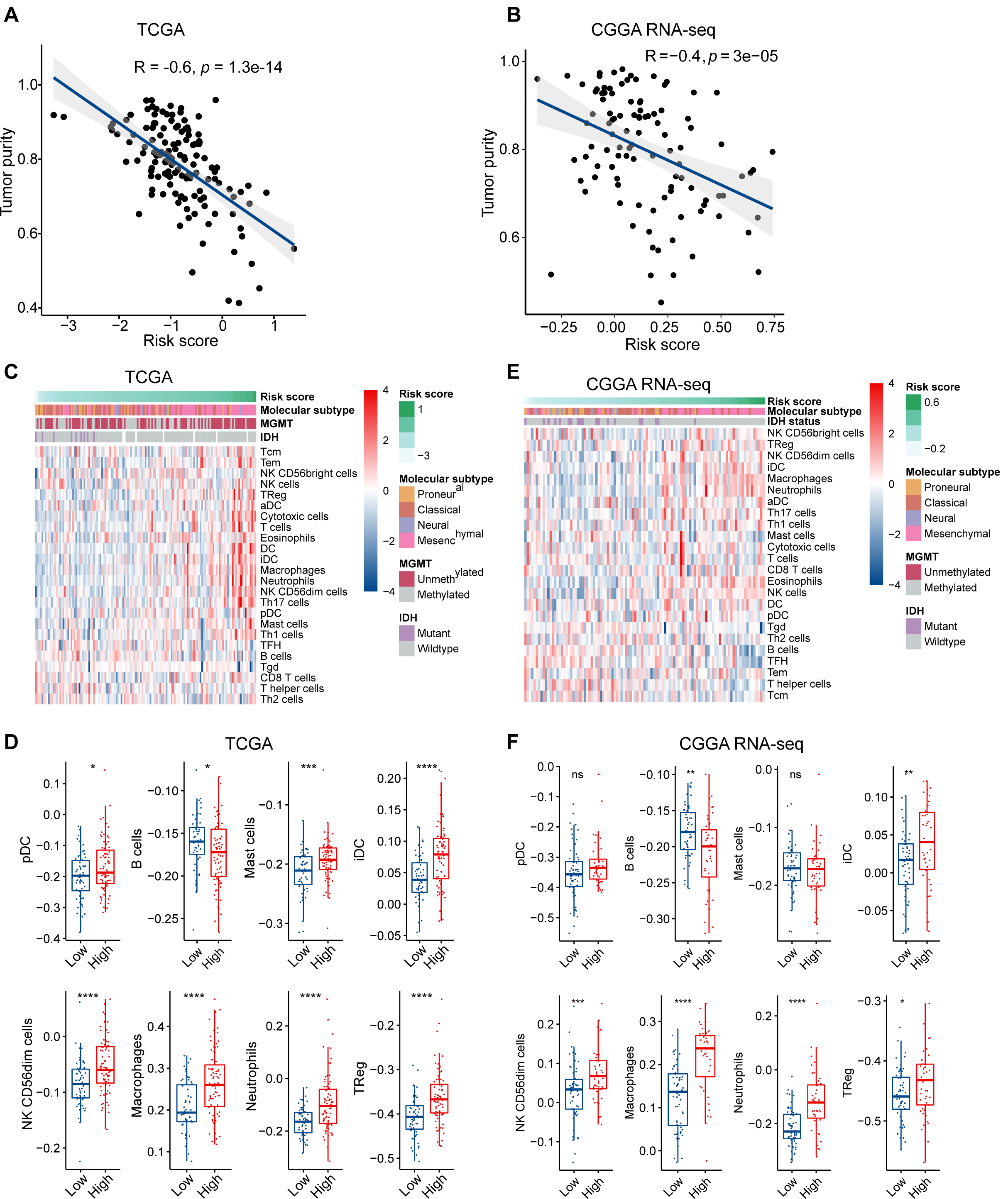
The risk signature is associated with tumor purity and immune infiltration in the microenvironment of GBM. Correlation of risk score and tumor purity in GBM samples in the TCGA (**A**) and CGGA RNA-seq (**B**) cohorts. The coefficient and P-value were obtained from Pearson correlation analysis. Heatmaps delineate the relationship between risk score and the contents of 24 types of immune cells infiltrated in the tumor tissues in the TCGA (**C**) and CGGA RNA-seq (**E**) cohorts. Box plots showing significantly different immune cells among high- and low-risk patients in the TCGA (**D**) and CGGA RNA-seq (**F**) datasets. * p < 0.05, ** p < 0.01, *** p < 0.001, **** p < 0.0001, with Mann-Whitney U test.

We then used the ssGSEA algorithm to evaluate the immune cell populations in the tumors from the two independent datasets. The contents of different types of immune cells varied as the risk score increased. The results showed that samples from the high-risk group in the TCGA cohort exhibited a higher abundance of pro-tumor immune cells, such as plasmacytoid dendritic cells (pDCs), immature dendritic cells (iDCs), CD56dim natural killer cells, macrophages, neutrophils, regulatory T cells (Tregs), and mast cells (P < 0.05) (Figure 4C, 4D). At the same time, B cells, which indicate a favorable prognosis of GBM patients (38), were highly abundant in the tumor samples from the low-risk group (P < 0.05). Similarly, high-risk patients in the CGGA RNA-seq cohort had significantly higher quantities of pro-tumor immune cells, except for pDCs and mast cells (P < 0.05) (Figure 4E, 4F). Therefore, we concluded that the immune-related biological processes and pathways associated with the risk score might result from the observed significant differences in differentiation or recruitment of various immune cell types.

### The risk score is correlated with immunosuppressive processes and the inflammatory responses

To investigate the potential mechanisms underlying the correlation between the genes related to the immune response and the risk score, we assessed the relationship between the risk score and glioma-associated immunosuppressors. The gene list of glioma-related immunosuppressors, including immunosuppressive cytokines and checkpoints, tumor-supportive macrophage chemotactic and skewing molecules, immunosuppressive signaling pathways, and immunosuppressors were obtained from the published literature (39) (40). The expression of immunosuppressive genes in tumors from the two GBM cohorts was plotted against the increasing risk score (Figure S5A, 5B). Following ssGSEA analysis, the score of each immunosuppression-related term was calculated for each tumor sample. Interestingly, the risk score was positively correlated with the identified immunosuppression-related terms in both the TCGA and CGGA RNA-seq cohorts (Figure 5A, 5B). This finding further corroborated that the immune gene signature is related to immunological suppression in GBM via changes in the expression of immunosuppressors.

**Figure 5.**
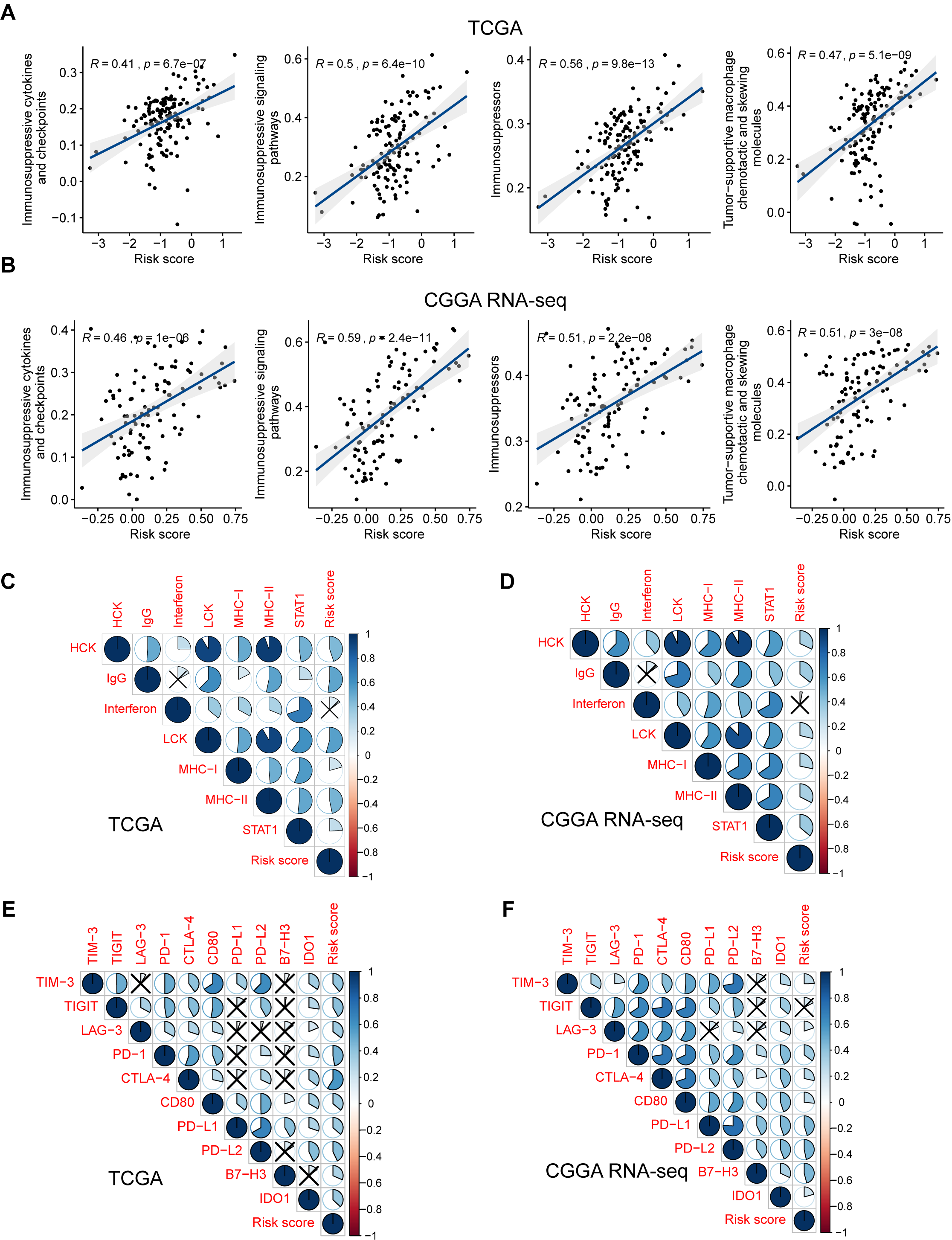
The risk signature is associated with immunosuppression and inflammatory activities in GBM. Scatterplots showing the correlation between risk score and each immunosuppressive activity in the TCGA (**A**) and CGGA RNA-seq (**B**) cohorts. The coefficients and P-values were calculated from Pearson correlation analyses. Corrplots showing the correlation between risk score and seven types of inflammatory activities in the TCGA (**C**) and CGGA RNA-seq (**D**) cohorts. The black cross indicates no statistical significance. Corrplots showing the correlation between risk score and the expression levels of several immune checkpoint molecules in the TCGA (**E**) and CGGA RNA-seq (**F**) cohorts. The black cross indicates no statistical significance.

**Figure 6.**
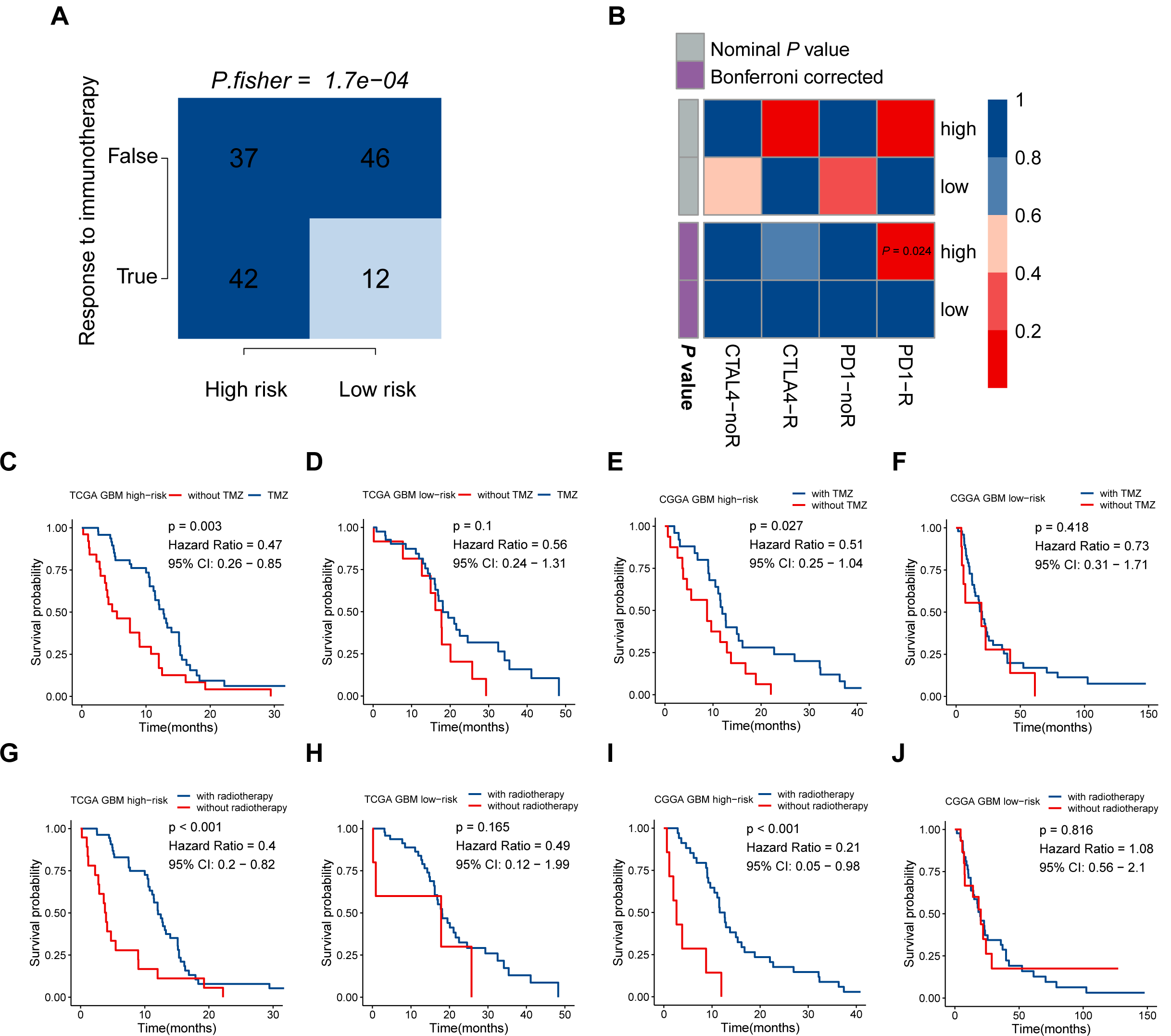
The risk signature may be tightly associated with the responses of immunotherapy, chemotherapy, and radiotherapy. **A**. Comparison of the predicted immunotherapeutic responses in different risk groups using the TIDE algorithm. P-value was obtained from Fisher exact test. **B**. The SubMap analysis concluded that high-risk patients might be more sensitive to anti-PD-1 immunotherapy. Patients from the high-risk group can benefit more from TMZ-based chemotherapy (**C, D**) and radiotherapy (**G, H**) in the TCGA GBM cohort, compared with those of the low-risk group. Similar responses to chemotherapy and radiotherapy were found in the CGGA RNA-seq GBM cohort (**E, F, I, J**).

As demonstrated above, genes correlated with the risk score were found to be significantly enriched in the category inflammatory response. Metagenes, i.e., expression patterns of various inflammation-related genes, were used to depict the relationship between the risk score and inflammatory activities in GBM. Except for interferon, most metagenes (HCK, IgG, LCK, MHC-I, MHC-II, and STAT1) were found to be positively associated with the risk score in both the TCGA and CGGA RNA-seq cohorts (P < 0.05) (Figure 5C, 5D, Figure S6A, 6B). These results revealed that the risk score was intimately tied to the B- and T-cell mediated immune response, as well as macrophage activation, but not to genes regulated by interferon during the remodeling of the immune microenvironment.

Pioneering investigations revealed that immunotherapy targeting immune checkpoints offered great hope for the clinical treatment of human cancers (41). We investigated the correlation between the risk score and the expression levels of several well-known immune checkpoints, both in the TCGA and CGGA RNA-seq datasets. Following Pearson correlation analysis, the risk score was found to be tightly associated with the expression of immune checkpoint molecules, including TIM-3, TIGIT (except in the CGGA RNA-seq cohort), LAG-3, PD-1, CTLA-4, CD80, PD-L1, PD-L2, B7-H3, and IDO1 (P < 0.05; Figure 5E, 5F). In conclusion, these findings implied a potential correlation of the risk score with the regulation of immunosuppression in GBM.

### High-risk patients are likely to be more sensitive to immunotherapy, chemotherapy, and radiotherapy

Given that the immune checkpoint blockade has not been used as a routine treatment for GBM, we modeled the potential response of patients in the different risk groups to immunotherapy, including antibodies against PD-1 and CTLA-4. Following the TIDE instructions, normal specimens from the GTEx database were used to adjust the gene expression matrix of GBM. The analysis indicated that the high-risk group (53.164%, 42/79) was significantly more likely to benefit from treatment with immune checkpoint inhibitors than the low-risk group (20.690%, 12/58) (P < 0.001) (Figure 6A, Table S8). SubMap is an unsupervised clustering method that can match subclasses in two independent gene expression data sets (29). Here, SubMap analysis further indicated that patients in the high-risk group are more likely to respond to anti-PD-1 immunotherapy (Bonferroni corrected P□= □0.024) (Figure 6B).

There is a large body of work implicating the influence of MGMT methylation status on the chemosensitivity to TMZ in GBM patients. Since the immune-gene-based signature was built based on the MGMT methylation status, we investigated whether this risk model can be used as an effective indicator for stratifying patients into different subgroups which may show distinct sensitivities to TMZ treatment. In both the TCGA and CGGA RNA-seq cohorts, whole GBM samples were divided into two groups according to their TMZ-based chemotherapy history, and then survival analyses for patients with high and low risk were carried out. In the TCGA cohort, patients who received TMZ treatment showed significantly better prognosis than those without chemotherapy in the high-risk group (HR = 0.47, 95% CI: 0.26-0.85, P = 0.003) (Figure 6C), while no significant difference was found in the low-risk group (P > 0.05) (Figure 6D). As we mentioned above, no difference in the proportion of MGMT methylation status between high- and low-risk groups suggested that the immune gene signature may serve as an effective biomarker for predicting the response of chemotherapy independent of MGMT methylation. The similar finding was also observed in the CGGA RNA-seq cohort that patients showed satisfactory responses to TMZ treatment in the high-risk group (HR = 0.51, 95% CI: 0.25-1.04, P = 0.027) (Figure 6E), while no difference was found between TMZ treated and non-TMZ treated subgroups in the low-risk group (P > 0.05) (Figure 6F). As for the role of this immune gene signature in regulating the sensitivity to radiotherapy in GBM patients, the same methods were performed as we described above. Patients with high risk can benefit more from radiotherapy both in the TCGA (HR = 0.4, 95% CI: 0.2-0.82, P < 0.001) and CGGA RNA-seq (HR = 0.21, 95% CI: 0.05-0.98, P < 0.001) cohorts (Figure 6G, 6I), while no difference existed in the low-risk group (Figure 6H, 6J). These findings indicated that the five-gene signature might serve as an effective biomarker for directing the applications of immunotherapy as well as routine treatment, including chemotherapy and radiotherapy.

### Development of a nomogram based on MGMT methylation-related signature

To develop a quantitative tool for predicting the prognosis of GBM patients in the TCGA cohort, we established a nomogram by integrating clinicopathological risk factors and immune gene signature based on the multivariable Cox proportional hazards model (Figure 7A). The point scale in the nomogram was utilized to generate point to these variables, and the risk of death of each GBM patient was qualified by accumulating total points of all variables. The risk score was found to have the most excellent weight among all these variables, which was consistent with the result of the previous multivariable Cox regression analysis. The C-index of this nomogram reached 0.704 (95% CI: 0.672-0.736). The development of the calibration plots further confirmed the significant consistency between predicted and observed actual clinical outcomes of GBM patients (Figure 7B). We additionally constructed a nomogram for the CGGA GBM cohort based on the risk model and other available clinicopathological features, and the feasibility of this nomogram was confirmed using the calibration plots (Figure 7C, 7D).

**Figure 7.**
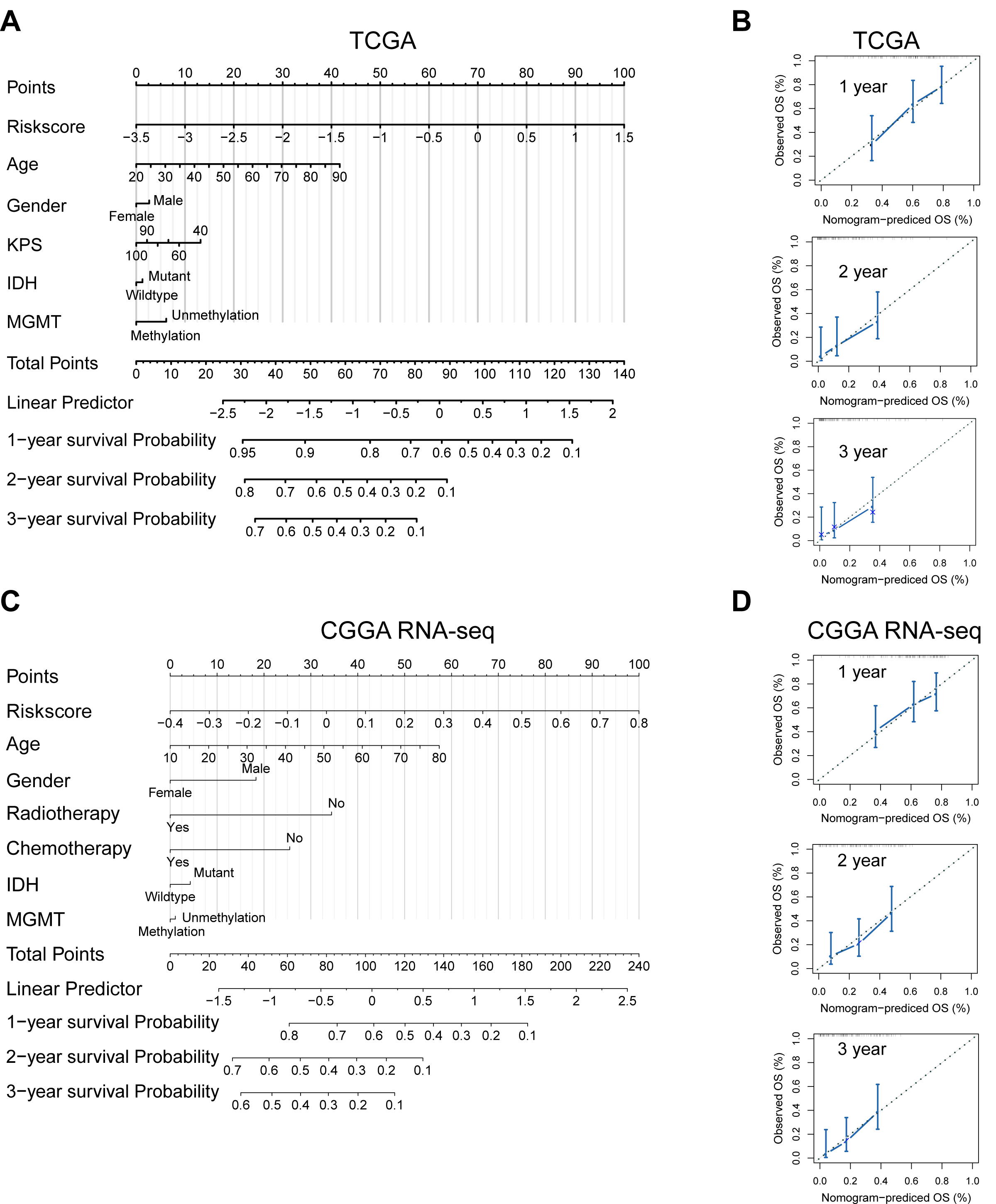
Developed nomogram to predict the probability of survival in GBM patients. Nomogram built with clinicopathological factors incorporated estimating 1-, 2- and 3-year overall survival for GBM patients in the TCGA (**A**) and CGGA RNA-seq (**C**) cohorts. The calibration curves describing the consistency between predicted and observed overall survival at different time points in the TCGA (**B**) and CGGA RNA-seq (**D**) cohorts. The estimated survival was plotted on the x-axes, and the actual outcomes were plotted on the y-axes. The gray 45-degree dotted line represents an ideal calibration mode.

## Discussion

In GBM, MGMT promoter methylation status is an essential determinant of the aggressiveness of the tumor. Malignant gliomas often show resistance to alkylating drugs due to the increased expression of MGMT, which remains a significant obstacle to clinical treatment (42). Previously, considerable efforts have been made to exploit the mechanisms of chemo-resistance induced by MGMT promoter demethylation. Over the past few years, the promising outcomes of immunotherapy in malignant tumors have attracted enormous attention. However, whether tumor immunity can be affected by the methylation status of the MGMT promoter has, to our best knowledge, not been explored before. Here, we sought to investigate the potential immunological mechanisms associated with the methylation status of MGMT in GBM by a comprehensive bioinformatic analysis. We addressed the critical role of MGMT methylation in regulating the remodeling of the tumor microenvironment of GBM.

The solid tumor mass of GBM is composed of both cancer cells and non-cancerous cells, such as immune cells, endothelial cells, fibroblasts, as well as some CNS-specific cell types such as astrocytes and microglia (43) (44). The discovery that multiple immune cell types can infiltrate the blood-brain barrier and the presence of reactivity against antigens derived from the CNS has made the view of the brain as an immunologically privileged site obsolete (45) (46). The most abundant immune cells, macrophages, comprise up to 30% of the total cells inside a glioma tumor (47). Tumor-associated macrophages suppress T cell responses due to insufficient expression of molecules associated with T cell co-stimulation(48). Dendritic cells are responsible for presenting antigens to T cells to induce immune responses in the CNS. The application of dendritic cell-based vaccination in GBM patients has produced survival benefits due to enhanced stimulated T-cell responses (49). Additionally, several studies demonstrated that reprogramming immunosuppressive T-cell subsets can stimulate antitumor immunity in glioma (48) (50). In this study, we discovered that genetic signatures of immune-related biological processes and signaling pathways were highly enriched in MGMT unmethylated specimens. Then, we filtered differentially expressed immune-related genes according to the MGMT promoter status and generated a signature based on six genes that can predict the clinical outcomes of GBM patients. At the same time, we investigated the contents of immune cells in the bulk tumor to predict immune infiltration using a marker gene-based approach based on transcriptomic data. Notably, we found that the infiltrating immune cell types differed significantly between the high and low-risk groups. In agreement with our hypothesis, the tumors in the high-risk group were severely infiltrated by immune cells characterized as belonging to pro-tumor subsets. Thus, this risk model provides an immunological interpretation of the effect of MGMT methylation status on the prognosis of GBM patients.

There is a large body of work that confirms the immunity-related roles of the genes identified in this study. For example, the transcription factor GATA3 has been characterized as a critical regulator in the development of T, B, NK, and innate-like lymphoid cells (51). GATA3 is responsible for maintaining the function of mature T and Treg cells via the IL-7 and IL-2 signaling pathways, respectively (52). Additionally, ectopic expression of GATA3 was found to promote the function of human type-2 innate lymphoid cells (53). TNSRSF9, also known as CD137, is a member of the TNF-receptor family that can activate cytokine production in cytotoxic T lymphocytes (CTLs) and Th1 cells by binding to its native ligand TNFSF9 (54). Recently, clinical trials with antibodies targeting CD137 have been launched with the expectation to improve cancer immunotherapy (55). VDR negatively regulates Forkhead box M1 (FOXM1) signaling and further suppresses the development of pancreatic ductal adenocarcinoma (56). In the immune system, VDR expression regulates the proliferation, differentiation, and function of CTLs (57). Stromal VDR activation reduces tumor-associated fibrosis and inhibits tumor-supportive signaling events in pancreatic cancer (58).

Immune suppression has been recognized as a hindrance to successful antitumor immunotherapy. Tumors create a microenvironment that sustains tumor growth as well as reduces adaptive immunity to tumor-derived antigens (59). We found a strong correlation between the risk score and immunosuppressive cytokines, tumor-supportive macrophages, as well as immunosuppressive signaling pathways. This finding implied that the risk score reflects the immunosuppressive microenvironment of GBM. Monoclonal antibodies that block immune checkpoint signaling have shown clinical benefits in several malignant tumors, including melanoma, NSCLC, Hodgkin lymphoma, and urothelial carcinoma. Such antibodies include pembrolizumab, nivolumab, and pidilizumab, which target PD-1, ipilimumab and tremelimumab for CTLA-4, or atezolizumab and durvalumab for PD-L1. Additionally, many clinical trials have been launched to evaluate the application of checkpoint inhibitors in GBM. A randomized phase III study (NCT02017717) was carried out to test the efficacy and safety of nivolumab alone versus bevacizumab (60). Clinical treatment of newly diagnosed GBM patients with ipilimumab and nivolumab is still under investigation (NCT02311920) (61). Therefore, we explored the relationship between the risk score and the expression levels of several well-known immune checkpoint molecules. The strong positive correlation implied that the identified signature mediates immune escape mechanisms in GBM, which provides a foundation for identifying novel therapeutic targets.

It is noteworthy that not all probands will respond to immune checkpoint blockade effectively in most cancer types (62). Various factors were reported as predictors for evaluating immunotherapy effectiveness, such as the mutation burden (63), tumor infiltration by cytotoxic T cells (64), PD-L1 expression (65), and tumor aneuploidy (66). The known mechanisms of tumor immune evasion include infiltration by CTLs and the prevention of T cell infiltration (67) (68). The TIDE method underscores these two distinct aspects and can predict the outcome of melanoma patients receiving immunotherapy more accurately than other factors (27). We demonstrated that more patients could benefit from immune checkpoint blockade in the high than in the low-risk group. Under a strict criterion of adjusted P-values, SubMap analysis further confirmed that anti-PD-1 therapy might produce favorable outcomes in high-risk patients.

## Conclusions

In summary, our study revealed that MGMT promoter methylation status is correlated with immunological processes and the remodeling of the tumor microenvironment in GBM. The developed immune-related gene signature could serve as a useful prognostic tool for GBM patients as well as for predicting which patients would benefit from immunotherapy. These findings may provide new insights into fundamental aspects of the critical role of MGMT methylation status in GBM.

## Acknowledgments

This work was supported by the National Natural Science Foundation of China under [grant number 81673210]. All authors would like to thank all contributors to the TCGA and CGGA project, especially.

---

**Table S1**. The detailed information about five GBM datasets used in the study.

**Table S2**. List of immune genes obtained from the ImmPort database and “GO_IMMUNE_RESPONSE” term.

**Table S3**. Gene enrichment in GBM patients with MGMT unmethylation.

**Table S4**. Differentially expressed genes between MGMT unmethylated and methylated samples in the TCGA GBM cohort.

**Table S5**. Multivariate Cox regression analysis of differentially expressed immune genes.

**Table S6**. Restricted mean survival (RMS) time ratio between low- and high-risk groups in different datasets.

**Table S7**. Genes strongly correlated with risk score in the TCGA and CGGA RNA-seq datasets.

**Table S8**. Predicted outcomes of immunotherapy for GBM patients (TIDE algorithm).

**Figure S1**. Correlations between the five prognostic immune genes in the TCGA (**A**) and CGGA RNA-seq (**B**) datasets.

**Figure S2**. A schematic view of MGMT methylation related immune gene selection and prognostic gene signature development.

**Figure S3**. Genes strongly associated with risk score in both the TCGA (**A**) and CGGA RNA-seq (**B**) cohorts. The correlation coefficients were obtained from Pearson correlation analyses.

**Figure S4**. Biological functions of the risk signature in the CGGA RNA-seq dataset. **A**. Bar plot showing the top 20 terms derived from the gene set enrichment of risk score. The x-axis represents statistical significance. **B**. The enrichment network plot visualizing the relationship between a set of representative terms. Each term is assigned with a unique color.

**Figure S5**. The risk signature is associated with immunosuppression in GBM samples. Heatmaps illustrating the association between risk score and glioma-associated immunosuppressive activities in the TCGA (**A**) and CGGA RNA-seq (**B**) cohorts.

**Figure S6**. The risk signature is associated with the inflammatory response in GBM samples. Heatmaps illustrating the association between risk score and predicted inflammatory activities in GBM samples from the TCGA (**A**) and CGGA RNA-seq (**B**) cohorts.

