## Supplementary material for "Molecular and clinicopathological characterization of a prognostic immune gene signature associated with MGMT methylation in glioblastoma": Figure S

**Supplementary figures**

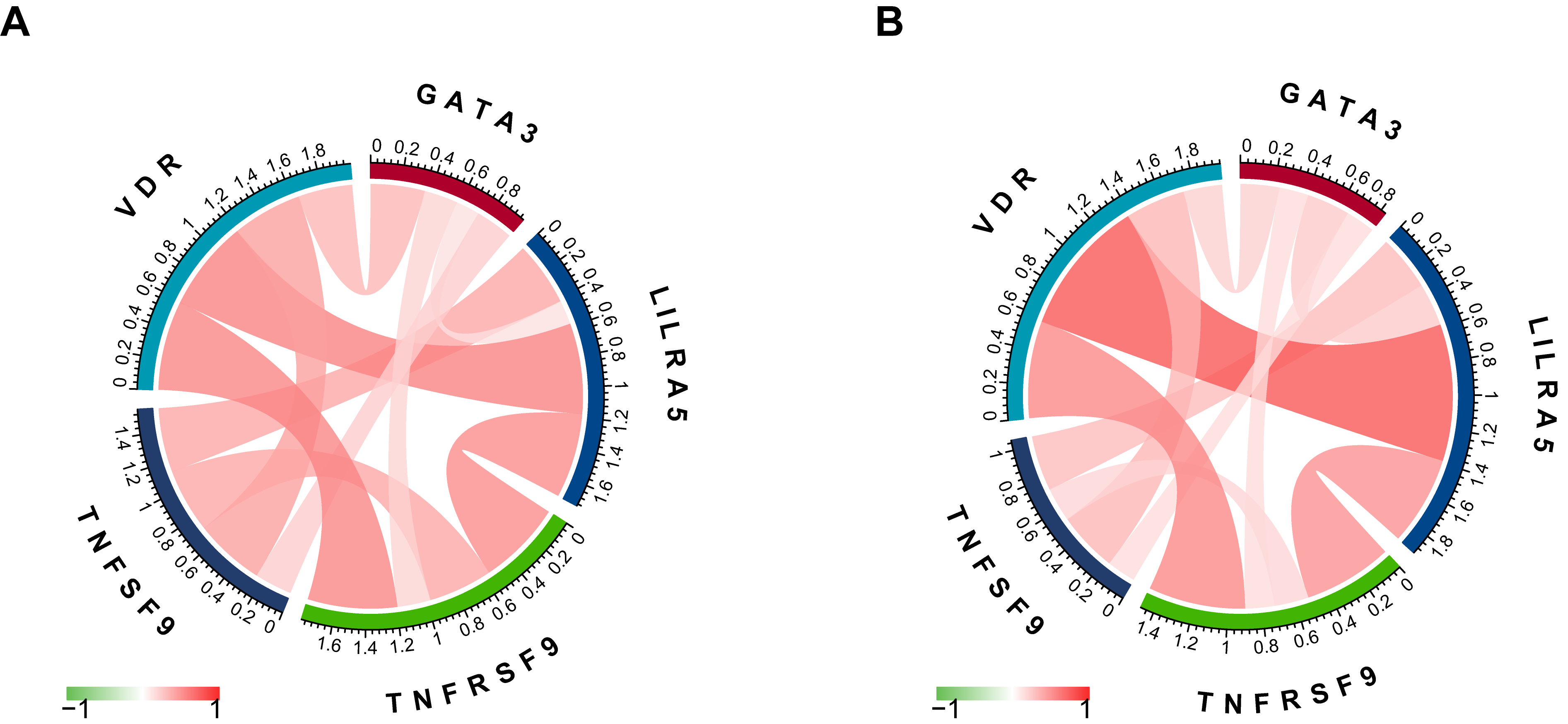

**Figure S1.** Correlations between the five prognostic immune genes in the TCGA (**A**) and CGGA RNA-seq (**B**) datasets.

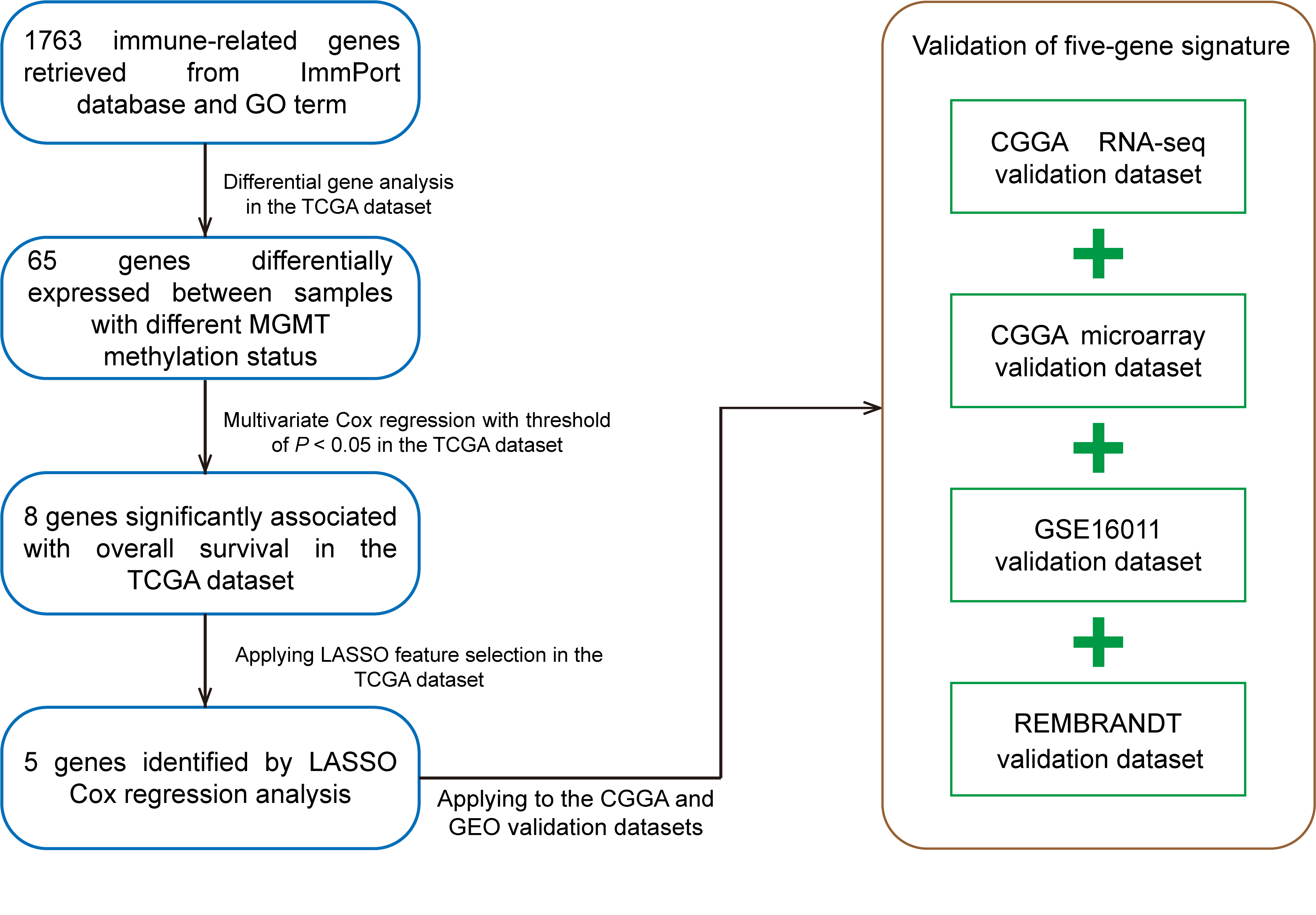

**Figure S2.** A schematic view of MGMT methylation related immune gene selection and prognostic gene signature development.

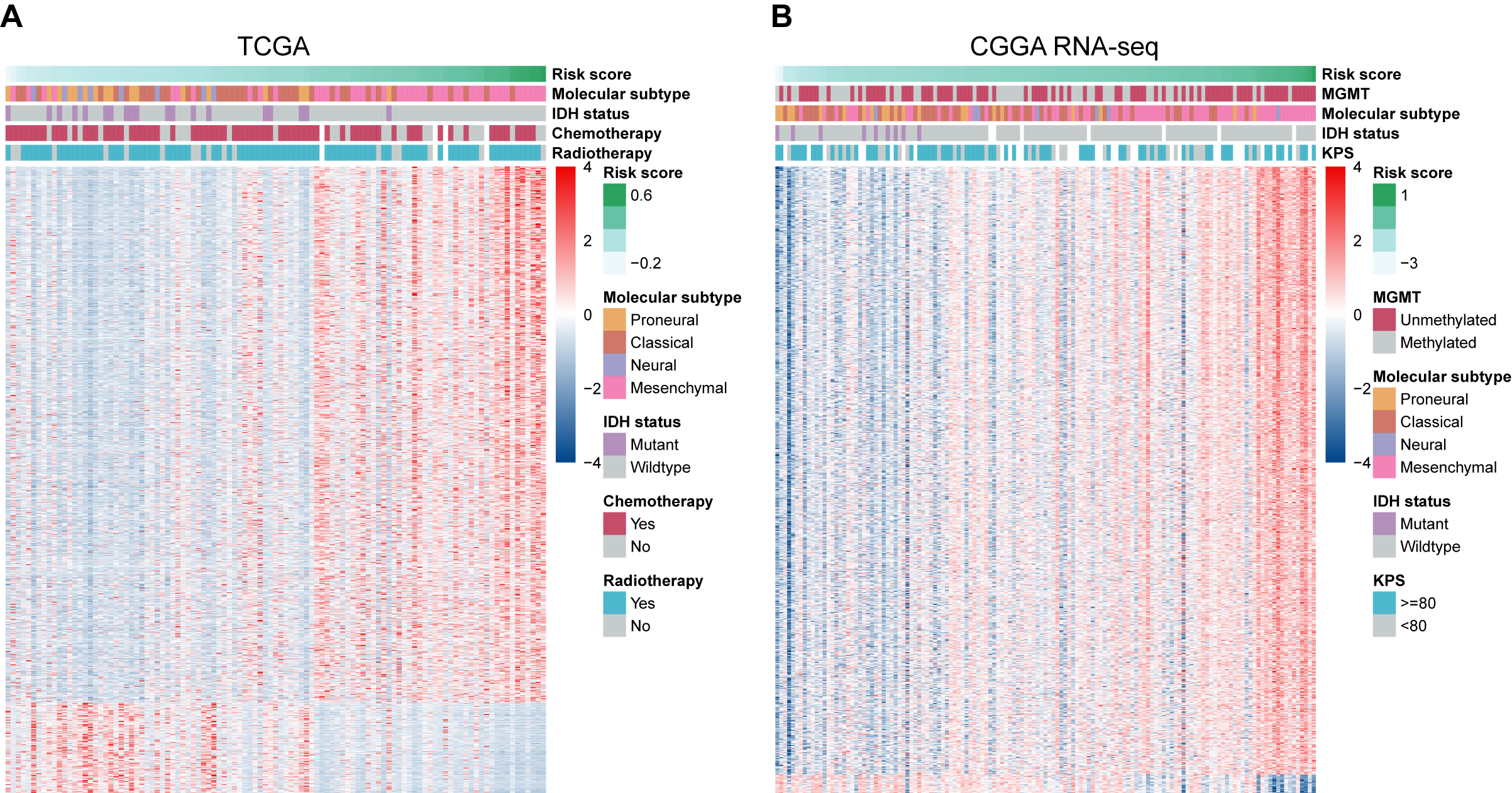

**Figure S3.** Genes strongly associated with risk score in both the TCGA (**A**) and CGGA RNA-seq (**B**) cohorts. The correlation coefficients were obtained from Pearson correlation analyses.

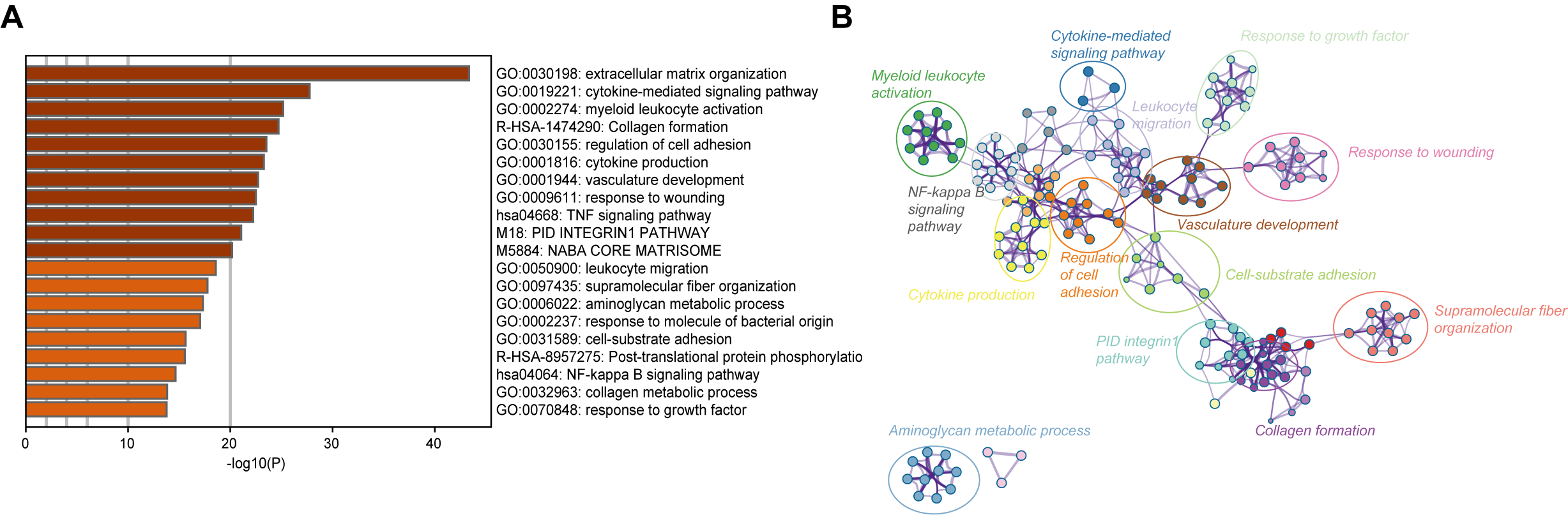

**Figure S4.** Biological functions of the risk signature in the CGGA RNA-seq dataset. **A.** Bar plot showing the top 20 terms derived from the gene set enrichment of risk score. The x-axis represents statistical significance. **B.** The enrichment network plot visualizing the relationship between a set of representative terms. Each term is assigned with a unique color.

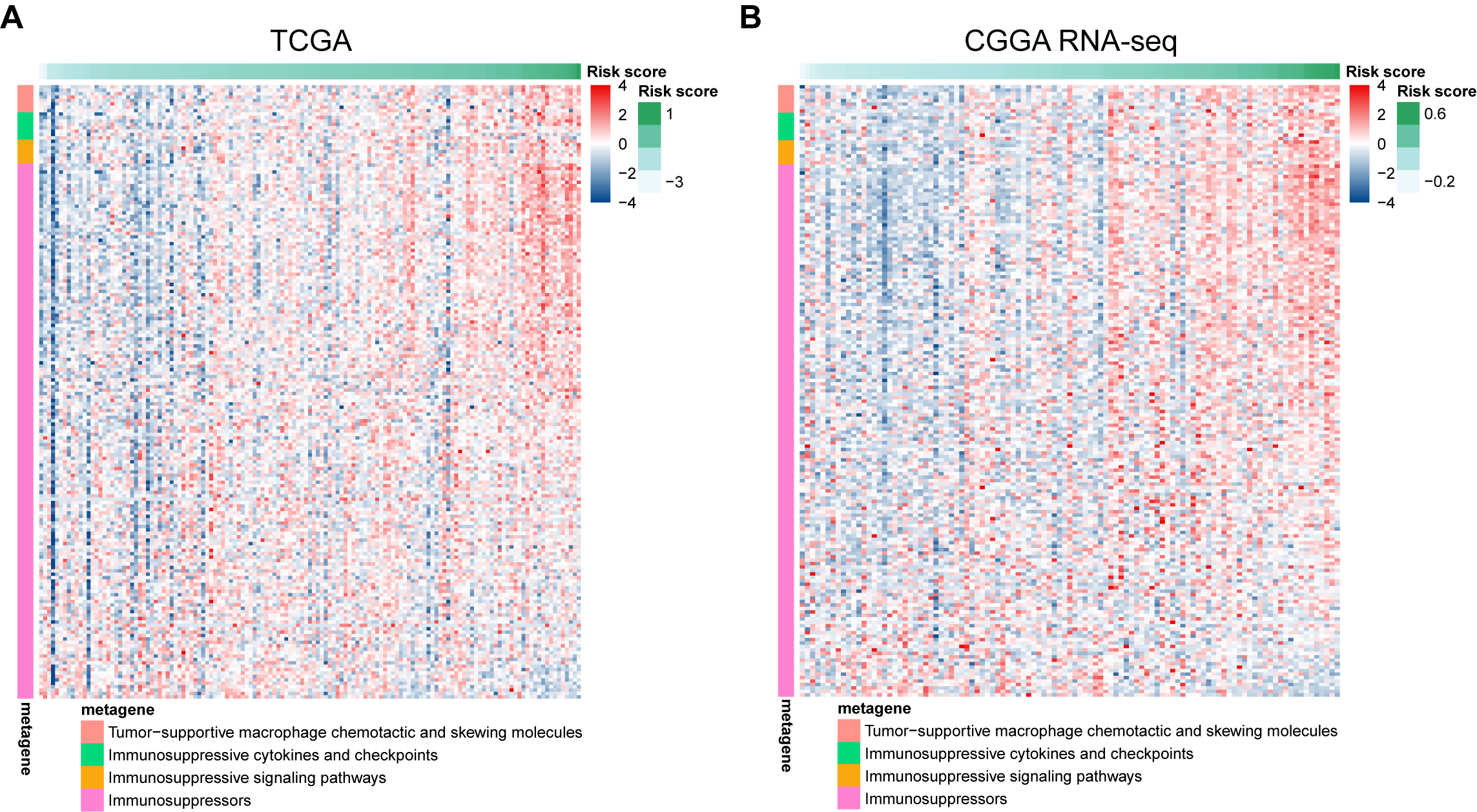

**Figure S5.** The risk signature is associated with immunosuppression in GBM samples. Heatmaps illustrating the association between risk score and glioma-associated immunosuppressive activities in the TCGA (**A**) and CGGA RNA-seq (**B**) cohorts.

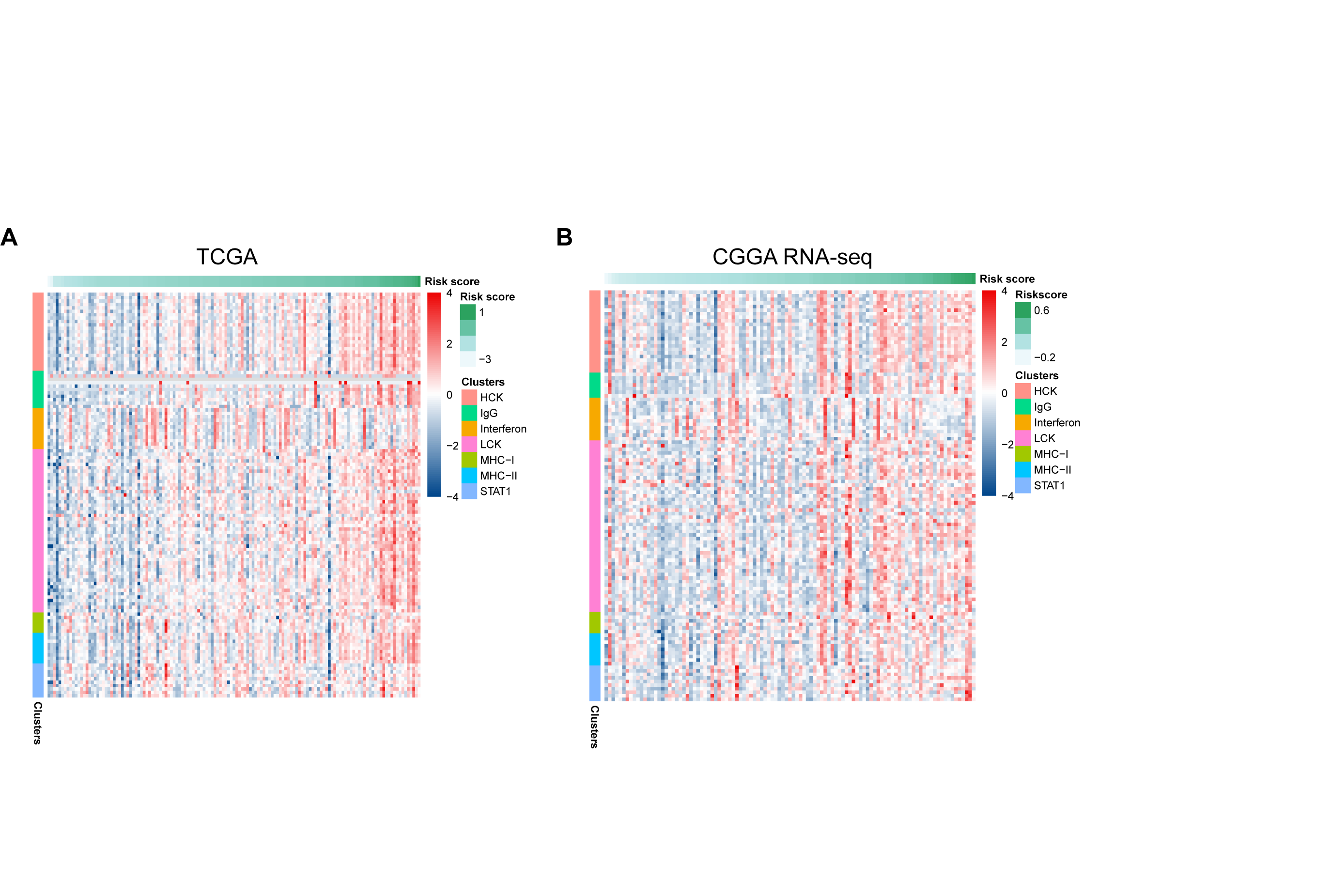

**Figure S6.** The risk signature is associated with the inflammatory response in GBM samples. Heatmaps illustrating the the association between risk score and predicted inﬂammatory activities in GBM samples from the TCGA (**A**) and CGGA RNA-seq (**B**) cohorts.
