## Supplementary material for "Molecular and clinicopathological characterization of a prognostic immune gene signature associated with MGMT methylation in glioblastoma": Table S1

| **GBM datasets** | **Number of samples** | **Platforms** | **Link** |
| --- | --- | --- | --- |
| TCGA | 165 | Illumina HiSeq | https://portal.gdc.cancer.gov/ |
| CGGA RNA-seq | 113 | Illumina HiSeq | http://www.cgga.org.cn/download.jsp |
| CGGA microarray | 112 | Agilent Whole Human Genome Array | http://www.cgga.org.cn/download.jsp |
| GSE16011 | 159 | Affymetrix GeneChip Human Genome U133 Plus 2.0 Array | https://www.ncbi.nlm.nih.gov/geo/query/acc.cgi?acc=GSE16011 |
| REMBRANDT | 220 | Affymetrix Human Genome U133 Plus 2.0 Array | https://www.ncbi.nlm.nih.gov/geo/query/acc.cgi?acc=GSE108474 |

**Table S1. The detailed information about five GBM datasets used in the study.**
